# Rapid host switching by Varroa destructor: implications for pathogen transmission and contact-based Varroa control

**DOI:** 10.64898/2026.08.19.745672

**Authors:** Joy Deng, Zachary Y. Huang

## Abstract

**BACKGROUND:** *Varroa destructor* is the major ectoparasite of honey bees and a vector of viral pathogens. Because pathogen transmission and exposure to contact-active acaricides depend on mite host contacts, understanding the factors governing host residence time is important for both disease epidemiology and pest management. We quantified host residence time under varying bee densities and host-type compositions.

**RESULTS:** Mean residence time was 9.48 h across 312 host-residence events. Mites remained on individual hosts for only 2.36 h on Day 1 but approximately 11–14 h from Day 2 onward. A generalized linear mixed model showed a strong positive effect of day on residence time (β = 0.341, SE = 0.044, P < 0.001), corresponding to an approximately 41% increase in residence time per day. Excluding Day 1 eliminated this effect (P = 0.16), indicating that the temporal pattern was driven primarily by the initial exposure period. Reconstructing Day 1 observations to an 8-hour schedule confirmed that this pattern was not an artifact of observation frequency. Neither host type nor bee density affected residence time, and mite occupancy of nurse bees matched host availability.

**CONCLUSION:** Host residence time was governed primarily by initial exposure rather than host identity or moderate crowding. The results identify a previously undescribed exploratory phase immediately after mites enter a novel adult-bee population. Because shorter residence times imply more frequent host switching, these findings improve our understanding of pathogen transmission dynamics and may help explain variation in the performance of contact-based *Varroa* control strategies.

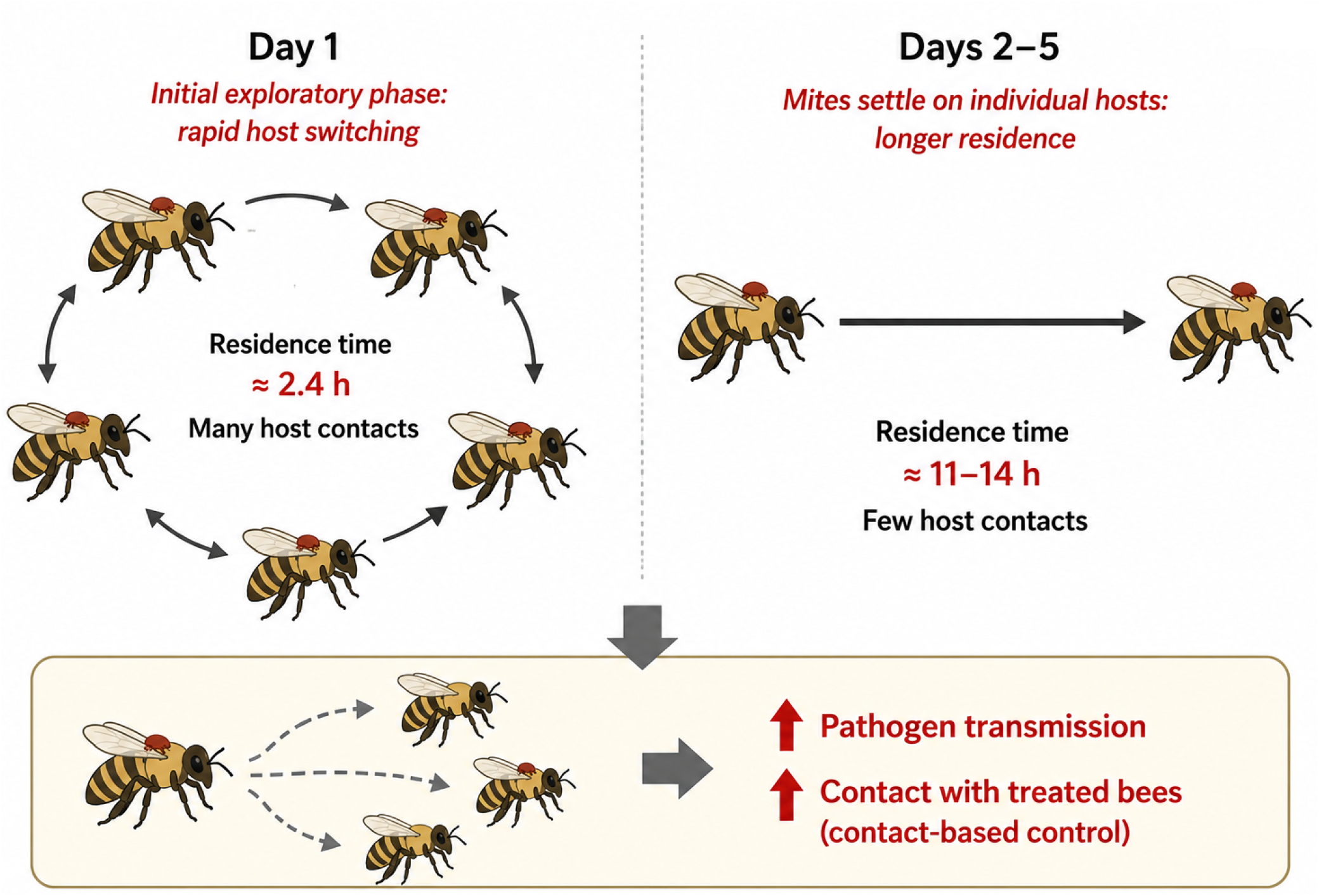

## Introduction

Honey bees (*Apis mellifera*) are critical pollinators in agricultural systems, yet managed colonies continue to experience substantial annual losses. In the United States, total colony mortality exceeded 55% in the 2024–2025 monitoring period [1]. A primary driver of these losses is the ectoparasitic mite *Varroa destructor*, widely recognized as the most significant biotic threat to honey bee health [2,3,4].

Beyond direct feeding damage, *Varroa* acts as a highly efficient vector of viral pathogens. It plays a central role in the amplification and transmission of deformed wing virus (DWV), a major contributor to colony collapse [5,6]. In addition, *Varroa* is associated with the transmission or increased prevalence of multiple other honey bee viruses, including acute bee paralysis virus (ABPV), Kashmir bee virus (KBV), Israeli acute paralysis virus (IAPV), sacbrood virus (SBV), black queen cell virus (BQCV), chronic bee paralysis virus (CBPV), and Lake Sinai viruses (LSV) [7,8]. Because viral spread depends on the number of adult hosts contacted by individual mites, understanding the factors governing host residence time—and consequently host-switching frequency—is central to predicting colony-level disease dynamics.

Host residence time may also influence the performance of contact-based Varroa control strategies. During the phoretic phase, many acaricides are acquired when mites contact treated adult bees. Because shorter residence times imply more frequent host switching, mites are expected to encounter treated workers more rapidly than mites remaining on a single bee for prolonged periods. Consequently, understanding the factors governing host residence time is important for both disease transmission and pest management.

In a recent study, Lamas et al. [9] demonstrated that promiscuous feeding across multiple adult hosts increases the vectorial capacity of *Varroa destructor*, estimating that mites remained on a single adult host for approximately 2.5 days on average. While this work provided key insights into transmission potential, it was conducted under simplified conditions with limited host diversity and relatively low worker densities compared to brood-nest environments. Moreover, little is known about how host residence time changes during the initial period after mites enter a new adult bee population.

Previous studies have shown that *Varroa destructor* do not distribute randomly among adult workers, with mites more frequently associated with nurse-age bees than with foragers [10,11]. In our previous work [12], we also observed a behavioral caste bias toward nurse-age workers under colony-relevant conditions. However, the mechanisms underlying this apparent preference remain unclear. The greater occupancy of nurse bees could arise from longer residence times once attached, or differences in encounter rates driven by spatial organization within the colony.

Here, we quantified host residence time as a direct measure of phoretic host use. Because shorter residence times necessarily result in more frequent host contacts over a given period, host-switching frequency can be inferred from residence time. Unlike previous studies in which mites were removed from their hosts before behavioral observations, mites in our experiments remained continuously attached to naturally infested newly emerged workers ("bee0") during introduction into the arena. This design avoided experimentally removing mites from their original hosts before observations began and allowed us to quantify their initial host residence time following entry into a novel social environment. We tested three predictions: (1) mites exhibit an initial exploratory phase characterized by unusually short host residence times; (2) higher density reduces residence time via increased encounter frequency; and (3) if nurse preference reflects active host selection, it should be reflected in either longer residence times on nurse bees or greater host occupancy of nurse bees during early exposure.

Because estimates of worker packing density in brood-nest regions are scarce, we additionally measured comb volume to approximate the spatial environment available to adult bees. This allowed us to compare the density of our experimental arenas with that expected under colony conditions.

## Materials and Methods

### Study organisms

Worker honey bees (*Apis mellifera*) were collected from healthy colonies and categorized into three functional groups based on behavioral role: nurse bees (bees with heads in larval cells), newly emerged workers (“newbees”), and foragers (bees returning to a colony with nectar or pollen). Approximately 30 individuals of each caste were collected per trial.

*Varroa destructor* mites were obtained from newly emerged (<24 h) adult workers from two highly infested colonies.

### Experimental arena and environmental conditions

Experiments were conducted in transparent Petri dishes. Either a cover or a bottom of a Petri dish was covered with a thin piece of Plexiglass, which had two openings: a small port fitted with a 3 mL syringe for delivering sugar solution and a larger opening for introducing bees. Dishes were maintained in an incubator (Thermo Scientific Forma Series II Water-Jacketed CO_2_ Incubator) at 34.0 ± 0.2 °C and 50 ± 5% relative humidity, approximating brood-nest conditions.

Bees were provided ad libitum access to 50% (w/v) sucrose solution via the feeding port, and food levels were checked daily to ensure continuous availability.

### Bee marking and preparation

To enable individual identification, bees were temporarily immobilized by chilling on ice after being placed in sealed plastic bags. Individuals were marked dorsally on the thorax using Testor”s paint. Up to eight unique markings were generated using combinations of four colors and two patterns (horizontal or vertical lines).

An unmarked mite-infested newly emerged worker (**bee0**) was introduced into each dish. Because this bee carried the focal mite at the start of the experiment, it was treated as a separate host category from the other newly emerged workers. Mite introductions typically occurred between 15:00 and 15:15, so the initial observation period (“Day 1”) lasted approximately 9 h, ending with the midnight observation. Thereafter, observations were conducted at regular 8-hour intervals.

### Experimental grouping and density manipulation

Marked bees were assigned to Petri dishes in mixed-caste groups. In Trials 1 and 2, each dish contained ten bees (three nurses, three newbees, three foragers, and one mite-infested bee0). In later trials, dishes contained four nurses, four foragers, and one bee0.

After introduction, each dish and the Plaxiglass were joined tightly with adhesive tape to prevent mite escape.

To test the possible effect of host density, two dish configurations were used: a lid (90.5 × 7 mm) and a base (86 × 12 mm), corresponding to volumes of approximately 45.0 mL and 69.7 mL, respectively. Five trials were conducted over a three-month period, each including 5–12 dishes. Bees were collected from five colonies, while mites were obtained from two separate source colonies.

### Observation protocol and behavioral recording

Each experimental group was observed for five days. Because mites were expected to switch hosts more frequently immediately after introduction, observations were more frequent on Day 1. On Day 1, dishes were checked at short intervals (10–20 min, depending on trial), followed by observations approximately 4 h after introduction and again at midnight, after which the regular 8-hour schedule began.

A total of 361 observations were recorded, yielding 312 host-residence events with measurable durations; in approximately 15% of observations, the host of a mite could not be determined.

At each observation, the identity of the host bee (because each has unique markings) was recorded for each mite and the host-type (bee0, nurse, forager, or newbee) was also recorded. The number of bees remaining in each dish was also recorded daily, with dead bees (if any) removed daily (and recorded).

Host residence duration was calculated as the elapsed time (hours) between successive observed host changes. Because Day 1 included shorter observation intervals, some residence durations were not multiples of 8 h. For example, a mite may have remained on a host for 3.75 h on Day 1 and subsequently for 16 h on Day 2, yielding a total residence duration of 19.75 h.

### Simulation of observation frequency

Because the initial observation period lasted only about 9 h before the regular 8-hour schedule began, all host-switching events occurring during this initial period were collapsed into a single apparent residence event for each mite, approximating the information that would have been obtained under an 8-hour observation schedule. These reconstructed Day 1 residence times were then compared with the actual observed residence times from Days 2–5, which had already been recorded at approximately 8-hour intervals.

### Estimation of worker packing density

Worker packing density was estimated by measuring comb volume using water displacement. Three frames were individually immersed in water, and displaced volume was quantified gravimetrically. Because some water entered open cells, these measurements underestimated true comb volume. Frame weights before and after immersion were used to correct for water absorption.

### Statistical analyses

All statistical analyses were done in R (version 4.5.2, R development, 2026) [13]. Host residence time (hours) was analyzed using generalized linear mixed models (GLMMs) with a Gamma error distribution and log link function implemented in the R package *glmmTMB*.

Fixed effects included day (treated as a continuous variable), host type, and bee density. Random intercepts were included for trial and mite identity to account for repeated measures and experimental structure.

Bee density was calculated for each observation as the number of living bees divided by the volume of the experimental arena (45.0 or 69.7 mL). Bee density was included as a continuous fixed effect in the Gamma GLMM.

To evaluate whether temporal patterns were driven by early observations, a second model excluding Day 1 data was fitted.

Host occupancy was evaluated by comparing the observed proportions of mite observations on foragers, nurses, and newly emerged workers with the expected proportions based on host availability at each observation. Expected proportions were recalculated throughout the experiment as bee mortality altered host composition.

Descriptive data are presented as means ± SE. For generalized linear mixed models, regression coefficients (β), their standard errors (SE), z values, and associated P values are reported.

## Results

### 1. Rapid exploratory phase following introduction of naturally infested newly emerged bees

Across all attachment events (N = 312), mean residence time on a host was 9.48 h. Residence time increased sharply after the initial exposure period (Fig. 1). On Day 1, mites remained on individual hosts for only 2.36 h on average, whereas from Day 2 onward mean residence time increased to approximately 11–14 h. A Gamma generalized linear mixed model revealed a significant positive effect of day on residence time (β = 0.341 (SE = 0.044), z = 7.79, P < 0.001), corresponding to an approximately 41% increase in expected residence time per additional day. However, when Day 1 observations were excluded, the effect of day was no longer significant (β = 0.037 (SE = 0.026), z = 1.40, P = 0.16), indicating that the temporal pattern was driven primarily by the contrast between the initial exposure period and subsequent days rather than by a continuous day-by-day increase.

**Fig. 1.**
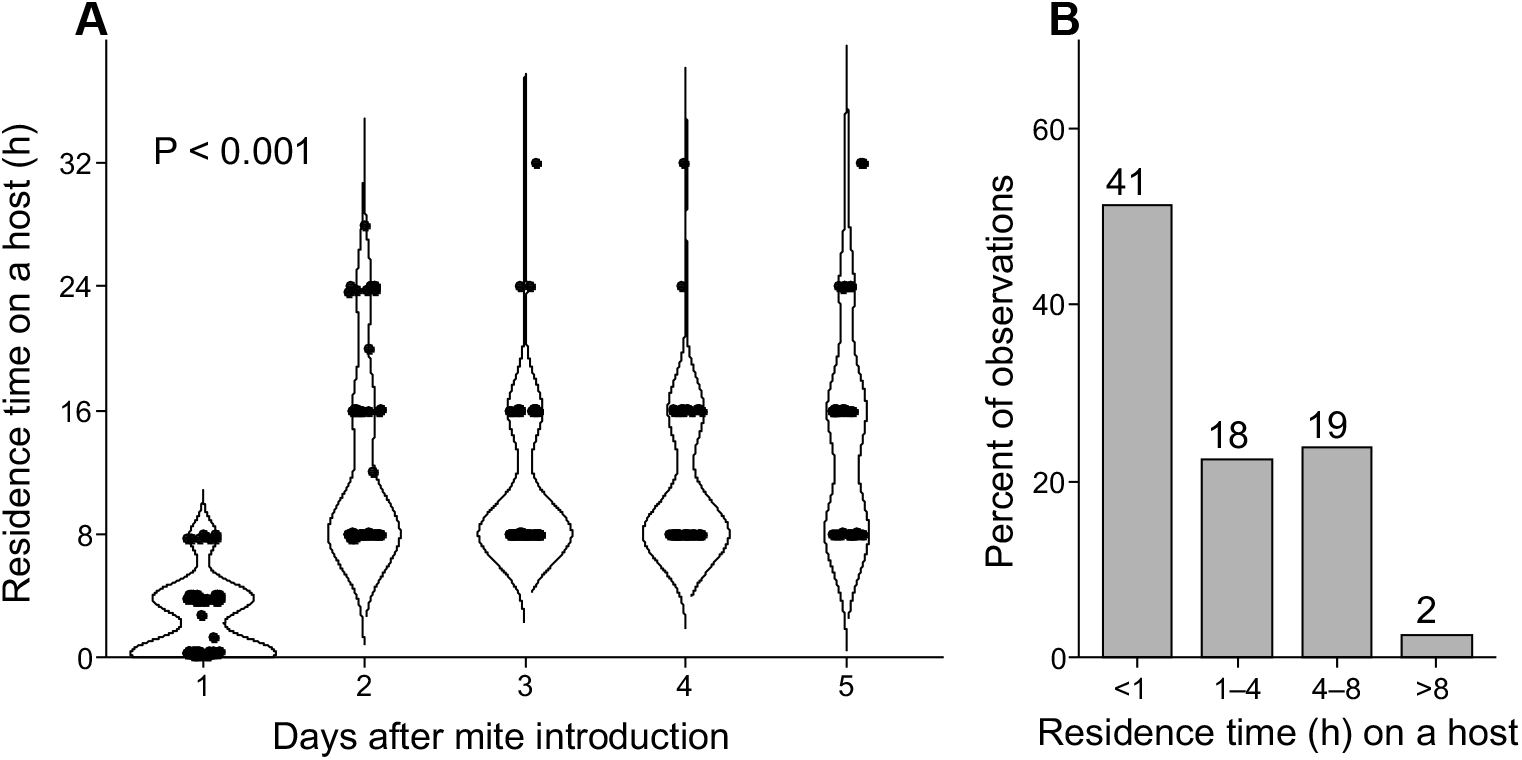
Residence time of *Varroa destructor* on individual honey bee hosts as a function of days after mite introduction (A). Distribution of Day 1 residence times (B), showing that more than 50% of observed residence events lasted less than 1 h.

To test whether this difference could be explained solely by observation frequency, all host-switching events occurring during the initial Day 1 observation period were collapsed into a single apparent 8-h residence event for each mite, thereby approximating the 8-hour observation schedule used from Day 2 onward. These reconstructed Day 1 values were then compared with the actual observed residence times from Days 2–5. Even under this conservative analysis, residence times remained significantly shorter on Day 1 than on subsequent days (Wilcoxon rank-sum test, W = 2940, P = 7.68 × 10−6; Fig. S1).

### 2. Bee density and host type did not affect residence time

Bee density ranged from 0.072 to 0.222 bees mL^−1^ and had no significant effect on host residence time (β = 0.492 (SE = 1.178), z = 0.42, P = 0.676; Fig. 2). Host type did not significantly affect residence time (all P ≥ 0.171 in the main model, Fig. 3), indicating that mites did not remain longer on any particular host type (newbee, nurse, or forager) under these confined conditions.

**Fig. 2.**
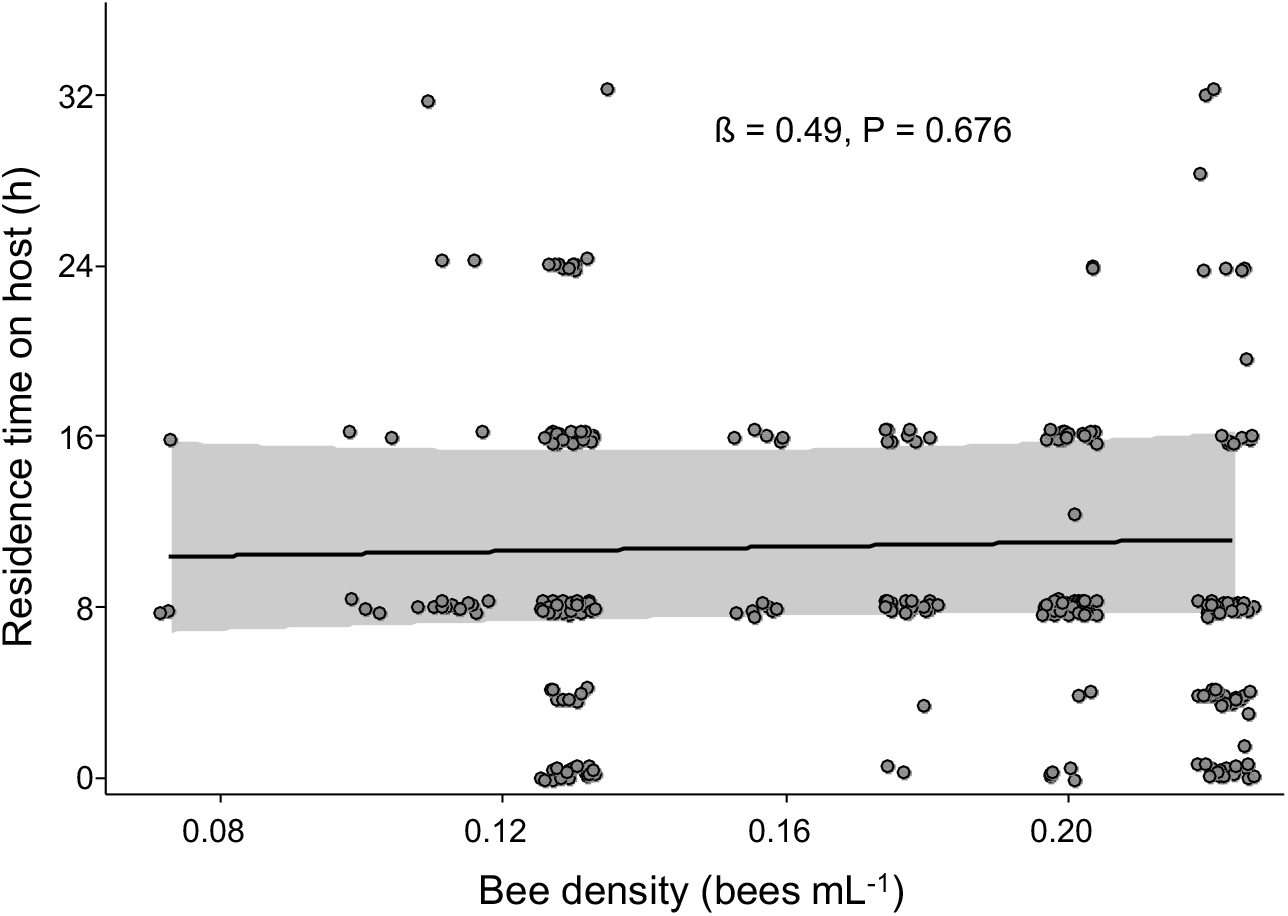
Relationship between bee density (bees mL^−1^) and *Varroa destructor* residence time. Each point represents one host residence event. The solid line shows the population-level prediction from the Gamma generalized linear mixed model, and the shaded region indicates the 95% confidence interval. Bee density had no significant effect on residence time (β = 0.49, P = 0.676).

**Fig. 3.**
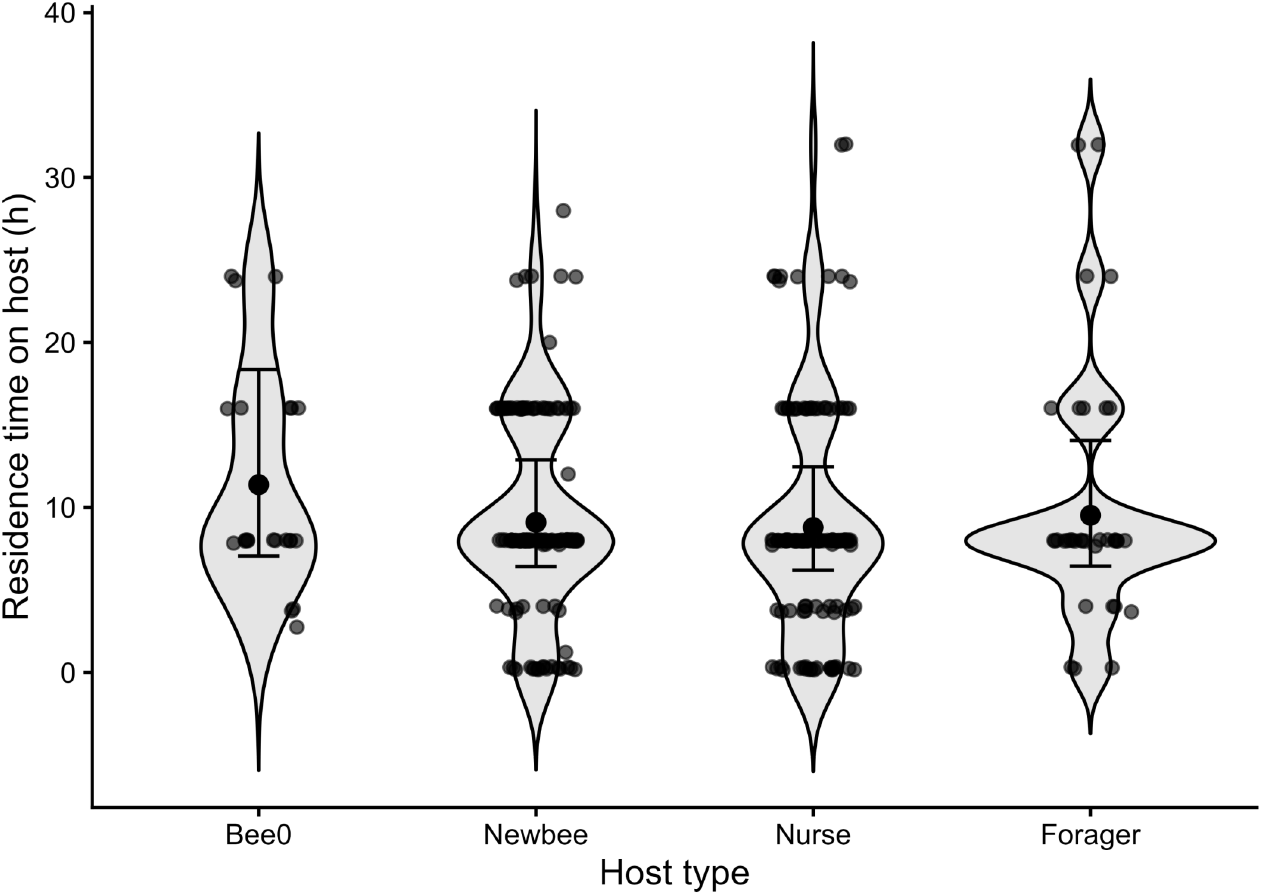
Residence time of *Varroa destructor* on individual honey bee hosts as a function of host type. Residence time did not differ significantly among host types (GLMM, P > 0.05). "Bee0" denotes the unmarked newly emerged worker carrying the mite when it was introduced into the experimental arena.

Overall host occupancy closely matched host availability (forager: 11.2% observed vs. 11.5% expected; nurse: 38.8% vs. 38.2%; newbee: 50.0% vs. 50.3%, P > 0.05).

### 3. Honey bee density in a hive

A deep frame was estimated to have a volume of 2.9 + 0.02 (mean ± SE) L. With 9–10 frames per hive body, the frames occupy approximately 20–25 L of space within a standard 42.5 L deep box. Assuming an average of 1,215 bees per side of a frame, this corresponds to approximately 21,870 bees in a fully occupied box. Based on these values, worker packing density is estimated at approximately 1.33 or 1.62 bees per mL (cm^3^), for a deep box with 9 or 10 frames, respectively. Having more frames in a colony reduces available space for workers, thus 10 frames caused a higher density for the same number of bees.

## Discussion

### Rapid exploratory phase following introduction of naturally infested newly emerged bees

The most striking result of this study is the strong temporal shift in host residence behavior. Mites exhibited very short residence times immediately after introduction, with Day 1 durations averaging only 2.36 h. From Day 2 onward, mean residence times increased to approximately 11–14 h. A Gamma GLMM detected a strong positive effect of Day across the full dataset; however, this effect disappeared when Day 1 observations were excluded, indicating that the overall pattern was driven primarily by the contrast between the initial exposure period and subsequent days rather than by a continuous day-by-day increase.

Because Day 1 observations were more frequent than those on subsequent days, shorter residence times could potentially reflect improved detection of brief residence events. However, this explanation was not supported. Even after reconstructing Day 1 observations under the same 8-hour observation schedule used from Day 2 onward, residence times remained significantly shorter (Fig. S1). This indicates that the observed difference reflects a genuine behavioral shift rather than an artifact of observation frequency.

Unlike previous studies, mites in our experiments were introduced while remaining attached to naturally infested newly emerged workers and were never detached or transferred immediately before behavioral observations. Consequently, the first host switch represented movement from an established host following entry into a novel social environment, rather than an experimentally induced host-searching phase following removal from the original host. This distinction may help explain why we observed a pronounced exploratory phase whereas Lamas et al. [9] reported substantially longer residence times from the beginning of their observations. In that study, mites were collected from phoretic adult bees, transferred to pupae, allowed to feed for two days before experiments began, and observations were conducted at longer time intervals. Differences in mite handling, prior feeding history, physiological state, and observation protocol may all have contributed to the contrasting patterns.

### Higher density does not affect residence time within the tested range

Contrary to our original expectations, host density did not significantly affect residence time within the range tested in this study. We found no density effect over the tested range (0.07–0.22 bees mL^-1^), but natural brood nests have much higher worker densities (1.3–1.6 bees mL^-1^), so density-dependent effects under colony conditions cannot be excluded. The much larger cup (16 oz = 473 mL) used by Lamas et al. [9] resulted in a bee density of only 0.017 bees mL^−1^, well below the range examined here. Our results therefore suggest that arena size alone was unlikely to explain the much longer residence times reported in that study.

### Host type has little effect on residence time

Contrary to expectations based on colony observations [12], host type did not significantly influence residence time. Mites did not remain longer on nurses than on other host types, and nurse bee occupancy closely matched host availability, indicating no evidence of preference for nurse bees under these conditions.

These findings suggest that, under confined arena conditions, switching behavior is governed primarily by host encounter rates rather than by strong host-type discrimination. Caste-based patterns observed in natural colonies may therefore arise from spatial organization or encounter structure rather than from active preference during individual switching events. Alternatively, "newbees" and nurses maintained in Petri dishes may rapidly acquire physiological characteristics of foragers in the absence of brood, comb and queen pheromones. Such changes can occur rapidly. For example, experimentally altering colony age demography reduced juvenile hormone titres within only 2.5 h [14]. Consequently, host-specific differences may have been diminished under our experimental conditions, making these results less representative than observations from intact colonies (Xie et al., 2016) [12].

### Implications for vectorial capacity

Previous work [9] demonstrated that promiscuous feeding across multiple adult hosts increases the vectorial capacity of *Varroa destructor*, emphasizing the epidemiological importance of switching frequency. Our results extend this framework by showing that host residence time is not constant through time but includes an initial exploratory phase during which mites remain on individual hosts for much shorter periods. Because shorter residence times necessarily result in more frequent host switching over a given period, this phase is expected to increase opportunities for host-to-host transmission. Consequently, estimates of vectorial capacity may depend not only on average residence time but also on the physiological or behavioral state of mites at the onset of the phoretic phase.

Across our arenas, mean residence times were approximately 2.36 h on the first day and about 11–14 h after the first day, both substantially shorter than the ∼2.5-day (about 60 h) estimate reported from the previous study [9]. That study did not use mites on newly emerged bees but mites from phoretic hosts. Furthermore, these mites were allowed to feed on pupae prior to the experiment. In addition, they used less frequent observation intervals (two times per day).

Because vectorial capacity depends directly on the number of host contacts per mite, shorter residence times imply increased opportunities for pathogen transmission. However, within the density range tested here, residence time did not vary detectably with host density, indicating that factors other than crowding—such as initial host exploration or prior feeding history—may play a more important role in determining switching rates under these conditions.

Host residence time may also have practical implications for Varroa control. Many contact-active acaricides rely on mites encountering treated adult workers during the phoretic phase. Mites that switch frequently among hosts would be expected to encounter treated bees more rapidly than mites exhibiting prolonged residence on individual workers, potentially influencing both the rate and uniformity of exposure to contact-active compounds. Although treatment efficacy was not examined here, our results suggest that variation in host residence time could influence both pathogen transmission and the performance of contact-based control strategies.

## Conclusion

Taken together, these findings indicate that host residence time is governed primarily by the initial exposure period rather than by host identity or moderate variation in bee density under the conditions tested. Residence time was shaped primarily by the initial exposure period rather than by host identity or moderate variation in density. Under confined conditions, *Varroa* host use appears to be opportunistic and encounter-driven rather than strongly selective.

These findings highlight the importance of considering behavioral context when interpreting host-switching rates and their epidemiological consequences. Phoretic residence time is not simply an intrinsic property of the mite, but a flexible outcome influenced by mite physiology and encounter structure. Future work comparing open and confined systems, or directly manipulating encounter rates while holding host composition constant, will help clarify how colony environment scales up to affect transmission dynamics.

## Acknowledgments

We thank Ling Chen for help during data collection and Dan Wyns for managing honey bee colonies used in this study.

## Figure legends

**Fig. S1.**
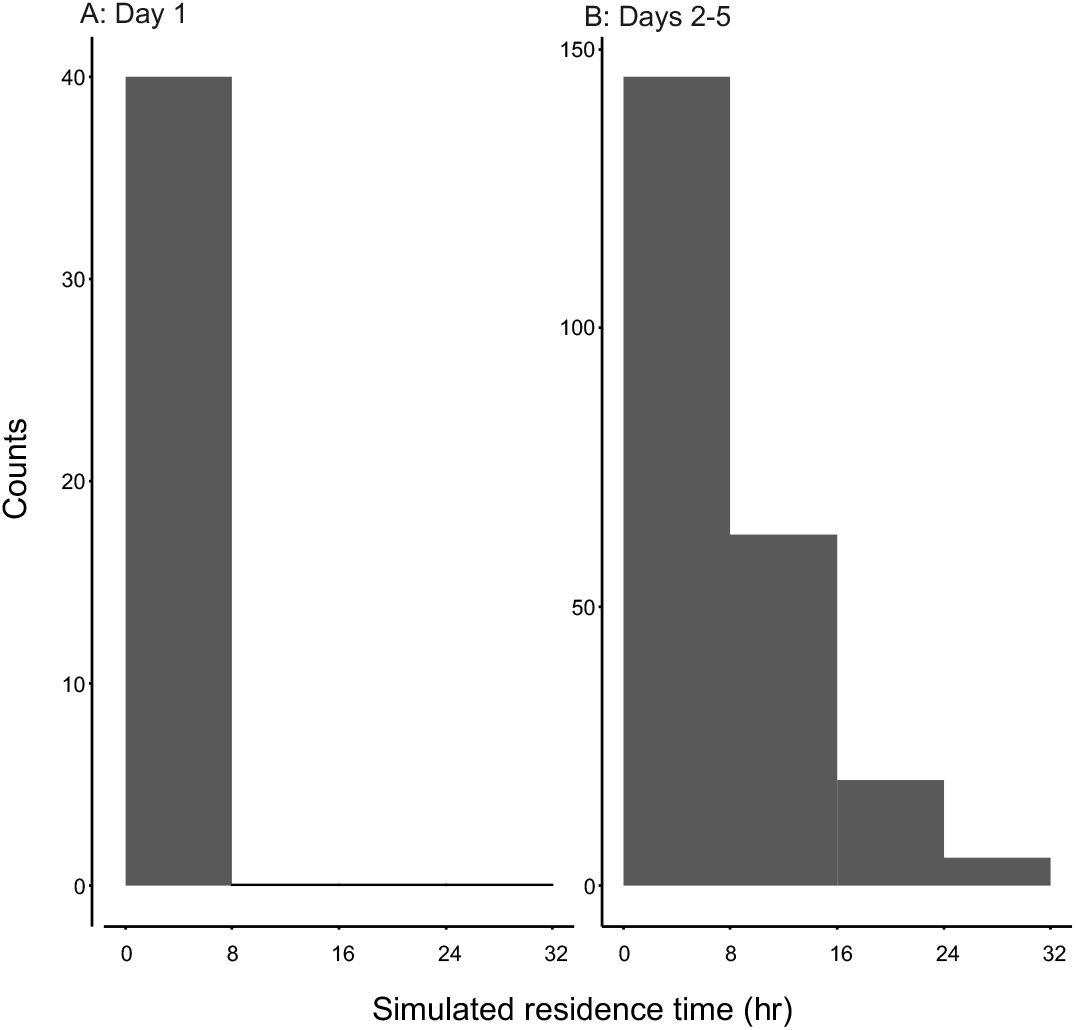
Simulated Day 1 residence times assuming an 8-hour observation schedule. (A) All host-switching events occurring before the first scheduled 8-hour observation on Day 1 were collapsed into a single apparent residence event for each mite (n = 40). (B) Actual observed residence times from Days 2–5, which were already recorded at approximately 8-hour intervals. Despite this conservative comparison, reconstructed Day 1 residence times remained significantly shorter than the observed residence times from Days 2–5 (Wilcoxon rank-sum test, W = 2940, P = 7.68 × 10−6).

